# Integrating prediction errors at two time scales permits rapid recalibration of speech sound categories

**DOI:** 10.1101/479089

**Authors:** Itsaso Olasagasti, Anne-Lise Giraud

## Abstract

Speech perception is assumed to arise from internal models of specific sensory features associated speech sounds. When these features change, the listener should recalibrate its internal model by appropriately weighing new versus old evidence in a volatility dependent manner. Models of speech recalibration have classically ignored volatility. Those that explicitly consider volatility have been designed to describe human behavior in tasks where sensory cues are associated with arbitrary experimenter-defined categories or rewards. In such settings, a model that maintains a single representation of the category but continuously adapts the learning rate works well. Using neurocomputational modelling we show that recalibration of existing “natural” categories is better described when sound categories are represented at different time scales. We illustrate our proposal by modeling the rapid recalibration of speech categories (Lüttke et al. 2016).

## Introduction

The way the brain processes sensory information to represent the perceived world is flexible and varies in response to changes in the stimulus landscape. Neurocognitive adaptation to varying stimuli can be driven by an external feedback signal, but might also take place with simple passive exposure to a changing environment. In the domain of speech perception, sound categories are susceptible to such stimulus-driven recalibration. Typically, unclear speech stimuli that are disambiguated by context (McQueen, Cutler, and Norris 2006; Clarke and Luce 2005) or by a concurrent stimulus presented e.g. in the visual modality (Bertelson, Vroomen, and De Gelder 2003; Vroomen et al. 2007), are subsequently perceived as a function of the disambiguated percept when presented again alone. Even simple exposure to novel statistics e.g., a variation in the spread of sensory features characteristic of stop consonants, quickly results in a modified slope in psychometric functions, and changes the way listeners classify stimuli (Clayards et al. 2008). Interestingly, acoustic perception is also changed by altered auditory feedback during production (Nasir and Ostry 2009; Lametti et al. 2014; Patri et al. 2018). These observations suggest that established speech sound categories remain largely plastic in adulthood.

Studies using a two-alternative forced choice task show that changes in acoustic speech categories can be induced by input from the visual modality (e.g. Bertelson, Vroomen, and De Gelder 2003). This effect is reciprocal; acoustic information can disambiguate lipreading (Baart and Vroomen 2010), resulting in measurable aftereffects. While such effects can be observed after repeated exposure to the adapting stimuli, recalibration can occur very rapidly (Vroomen et al. 2007). This fast and dynamic process has been modelled as incremental Bayesian updating (Kleinschmidt and Jaeger 2015) where internal perceptual categories are modified to better match the stimulus statistics. In their Bayesian approach, when listeners are confronted with modified versions of known speech categories, they alter the perceived category’s representation such as to make it more consistent with the actual features presented in the stimulus. The resulting recalibration weighs all evidence equally, independent of their recency. This model successfully describes perceptual changes observed when listeners are confronted with the repeated presentation of a single modified version of a speech sound. However, it cannot appropriately deal with intrinsically changing environments, in which the relevance of sensory cues decreases with time, with a faster decrease the more volatile the environment.

Inference in volatile environments has been studied mostly in relation to decision making tasks in which participants use an explicit feedback to keep track of the varying statistics of arbitrary cue-reward associations (e.g. Behrens et al. 2007) or arbitrarily defined categories (e.g. Summerfield et al. 2011). This shows that humans are able to adjust their learning rate to the volatility in the stimulus set, showing faster learning rates in more volatile environments, and vice versa, with faster learning rates translating into a stronger devaluation of past evidence. This has lead to normative models in which task relevant features are represented at a single yet variable time scale, with the focus of the models being on the online estimation of volatility (Behrens et al. 2007; C. Mathys et al. 2011).

Similar variable learning rates might apply to speech processing. However, we would like to propose that when recalibrating existing “natural” categories, a normative model should account for the fact that these categories may change at different time scales. Transient changes in how a speech category sounds, such as those coming from a new speaker, should not interfere with long-term or more slowly varying representations of the same category that perhaps encode commonalities across speakers. We therefore propose that speech sound categories should be represented by more than a single varying timescale.

Although speech category recalibration has not been systematically studied in variable environments, Lüttke and collaborators (Lüttke et al. 2016) found recalibration in an experiment that included audio-visual McGurk stimuli shown without an explicit adapting condition, and even for acoustic stimuli that were not ambiguous. In their study, which involved six different vowel/consonant/vowel stimuli presented in random order, the fact that the acoustic stimulus /aba/ was mostly perceived as an illusory /ada/ when presented with the video of a speaker producing /aga/ (McGurk effect), was powerful enough to yield observable adaptive effects across consecutive trials; specifically, the probability of an acoustic /aba/ to be categorized as /ada/ rather than /aba/, was higher when the trial was preceded by an audio-visual McGurk fusion. Lüttke, Pérez-Bellido, and de Lange (2018) further characterized the effect, from a signal detection theory perspective, as a change in how /aba/ sounds are processed. Other studies show that similar recalibration effects did not generalized across phonetic contrasts/categories. After participants had recalibrated acoustic sounds in the /aba/-/ada/ continuum (in which formant transitions are the main acoustic cue), recalibration was not present for /ibi/-/idi/ (main acoustic cue burst and frication), /ama/-/ana/ (also cued by formant transitions) or to /ubu/-/udu/ (same phoneme contrast also relying on formant transitions (Reinisch et al. 2014). Likewise, in a word recognition task participants were able to keep different F0/VOT correlation statistics for different places of articulation (Idemaru and Holt 2014). We hence hypothesized that such short-term changes modify the internal mappings between sensory features and the sublexical speech categories.

To test our hypothesis we simulated Lüttke et al.’s (Lüttke et al. 2016) experiment, using a model of audiovisual integration based on hierarchical Bayesian inference. We explicitly included a model for the McGurk effect based on our previous work (Olasagasti, Bouton, and Giraud 2015) and supplemented it with an adaptation mechanism that uses residual prediction error to update the internal representation – in the form of a hierarchical generative model-associated with the perceived category. The model divides the process in two steps; the first one is an on-line (moment-by-moment) prediction error minimization during the perceptual inference process, at the end of which there might remain “residual” prediction errors. This is typically the case in fused McGurk stimuli, since the best explanation for the multisensory input, (/ada/), is neither the audio /aba/ nor the visual /aga/, which leaves “residual” prediction errors in both acoustic and visual dimensions.

The proposed model was able to reproduce the results described above (Lüttke et al. 2016) only when we considered that adaptation occurred at two different time scales resulting in 1) a transient effect leading to recalibration towards the most recently presented stimulus features, which decayed towards 2) a longer-lasting representation corresponding to a mapping between category and stimulus features determined within a longer time span. Interestingly, a cascade model of synaptic modification seems to share some of these features (Iigaya 2016).

Overall, these findings are consistent with theories that posit that the brain uses generative – forward – models that are continuously being “recalibrated” to maintain self-consistency (e.g., K J Friston et al. 2010).

## Results

### Simulation of recalibration

The McGurk effect (McGurk and MacDonald 1976) offers a unique opportunity to assess cross-modality effects in adaptation. Lüttke et al. (Lüttke et al. 2016) in a re-assessment of an existing dataset selected participants with high percentage of fused percepts for McGurk stimuli. When presented with acoustic /aba/ together with a video of a speaker articulating /aga/ the most frequent percept was /ada/; we refer to these as fused McGurk trials. These listeners were combining information from the acoustic (A) and visual (V) modalities since the same /aba/ acoustic token was correctly categorized as /aba/ when presented alone. After a fused McGurk trial participants classified acoustic only /aba/ stimuli as /ada/ more frequently (29% /ada/ percepts) than when the acoustic only /aba/ was presented after any other stimulus (16% /ada/ percepts).

Our goal was to assess a model potentially showing how this effect fits within an ideal observer approach, using an internal generative model that interprets sensory input by continuously adapting to best match the incoming input. The model characterizes the sensory input by a visual feature, the amplitude of lip closure in the transition between the two vowels, which we denote by C_V_; and an acoustic feature, the amplitude of the 2^nd^ formant transition, which we denote by C_A_.

The internal generative model, described in detail in the methods section, generates continuous time lip motion and formant transitions for the congruent versions of each of the three possible tokens (k = /aba/, /ada/, /aga/). For our present purposes, the most important part of the model is its representation of the three congruent tokens. Each is represented by a Gaussian distribution in the two-dimensional feature space, itself a product of two unimodal Gaussian distributions centred at (θ_k,V_ θ_k,A_) and with standard deviations (σ_k,V_, σ_k,A_) for tokens k = {/aba/, /ada/, /aga/}. It assumes that given a speech token k, the C_A_ and C_V_ for each individual trial are chosen from the corresponding Gaussian distribution (Fig 1A). Once values for C_A_ and C_V_ have been determined, the unfolding in time of the lip closure and 2^nd^ formant transitions are obtained by multiplying a fixed temporal profile for each sensory feature (Fig 1B). During inference, the model is inverted and provides the posterior probability of a token given noisy sensory input.

**Fig 1.**
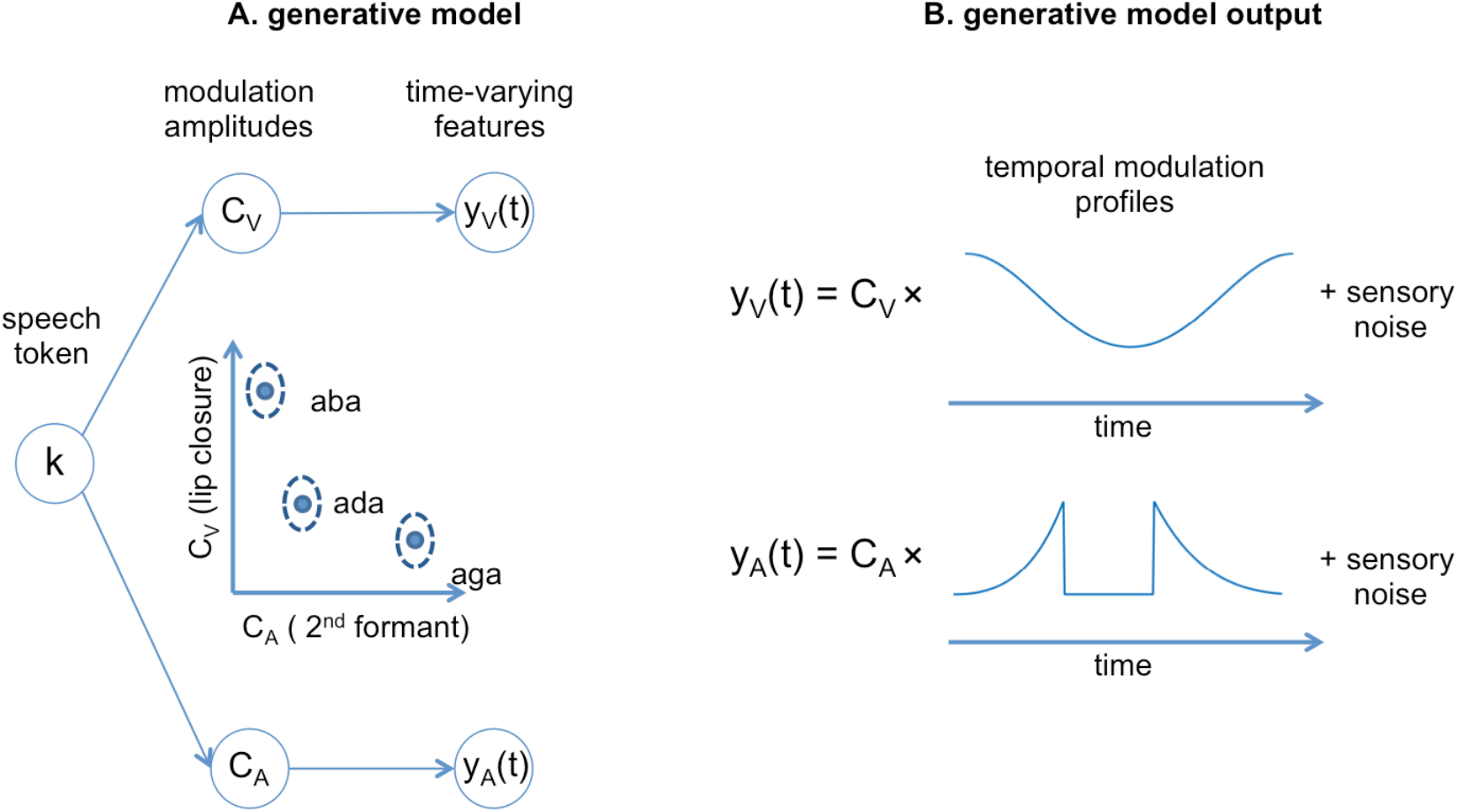
Schematics of the generative model. A. The schematic on the left shows the generative model. In a given trial, a speech token “k” determines the amplitudes of degree of lip closure (C_V_) and magnitude of second formant deflection (C_A_) by sampling from the appropriate Gaussian distribution. The distributions corresponding to each “k” are represented in the two-dimensional feature space in the middle. These values determine the amplitude of the time varying audiovisual features. B. The right panels show in more detail how the time varying output of the generative model is determined given a pair (C_V_, C_A_). The temporal profiles are fixed in the present model and are common for the three tokens, which are assumed to change only in their magnitude. The model also includes the “injection” of sensory noise. The lip modulation profile shows the lip movement in the transition between the two vowels. Similarly, the 2^nd^ formant transition profile shows the transition in the second formant between the two vowels.

The model qualitatively reproduces the average performance across participants in the task; we chose parameters that elicit a very high frequency of McGurk percepts (Fig 2B, middle panel of the bottom row) and assumed that listeners were always integrating the two sensory streams.

**Fig 2.**
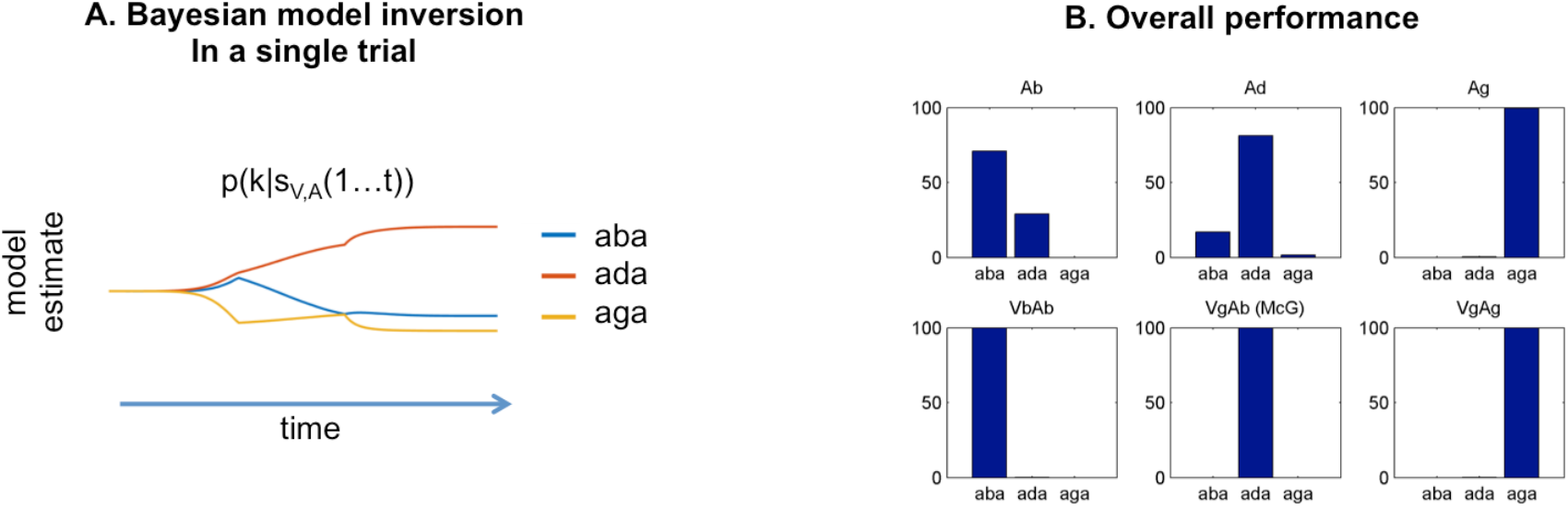
Model’s overall performance. A: Stimuli are presented to the simulated participant, which inverts the generative model and gathers evidence for each of the three possible outcomes. The example corresponds to a McGurk trial (acoustic /aba/, visual /aga/). The model’s posterior is maximum for /ada/. B: Simulation of the Lüttke et al. (2016) experiment. Model classification across all trials. Each subpanel shows the percentage of /aba/, /ada/ and /aga/ percepts corresponding to each of the six conditions (Ab: acoustic only /aba/; Ad: acoustic only /ada/; Ag: acoustic only /aga/; VbAb: congruent audivisual /aba/; VgAb incongruent McGurk stimuli with visual /aga/ and acoustic /aba/; VgAg: congruent /aga/). Congruent and acoustic only stimuli are categorized with a high degree of accuracy and McGurk trials are consistently fused, that is, perceived as /ada/. We reproduce the experimental paradigm consisting of six types of stimuli presented in pseudo-random order; three non-ambiguous acoustic only tokens: /aba/, /ada/ and /aga/, and three audiovisual stimuli: congruent /aba/, incongruent visual /aga/ with acoustic /aba/ (McGurk stimuli), and congruent /ada/.

In both acoustic and audiovisual trials listeners were asked to report the perceived acoustic stimulus in a three-alternative forced choice task. To simulate the listener’s choice, we calculated the probability of the acoustic token given the stimulus. In our notation, p(k_A_|s_V_,s_A_) for audiovisual stimuli and p(k_A_|s_A_) for unimodal acoustic stimuli, where s_A_ and s_V_ correspond to the sensory derived modulation amplitudes (s_A_ = y_A_/(acoustic modulation profile); s_V_ = y_V_/(lip modulation profile), see Materials and Methods for details). The percept at a single trial was determined by choosing the category that maximized the posterior. Fig 2A shows the model’s response to a McGurk stimulus; at the end of the trial the most likely interpretation corresponds to the syllable /ada/.

To model the recalibration after each trial, we changed the generative model’s location parameters associated with the perceived category. This was done for both the acoustic and visual parameters after an audio-visual trial, and for the acoustic parameter after an acoustic trial. We assumed that this happens for every trial and is part of the normal perceptual process. The changes are thus driven by any residual sensory prediction error.

When listeners consistently reported the fused percept /ada/ when confronted with a video of /aga/ and the sound of /aba/, the presence of the visual stream modified the acoustic percept from /aba/ to /ada/. Given that the acoustic input did not correspond to the one that was most expected from the perceived token, there was a systematic residual sensory prediction error. Since this residual prediction error was used as a signal to drive the model’s adaptation, the /ada/ representation moved towards the McGurk stimulus parameters after a fused percept (Fig 3A).

**Fig 3.**
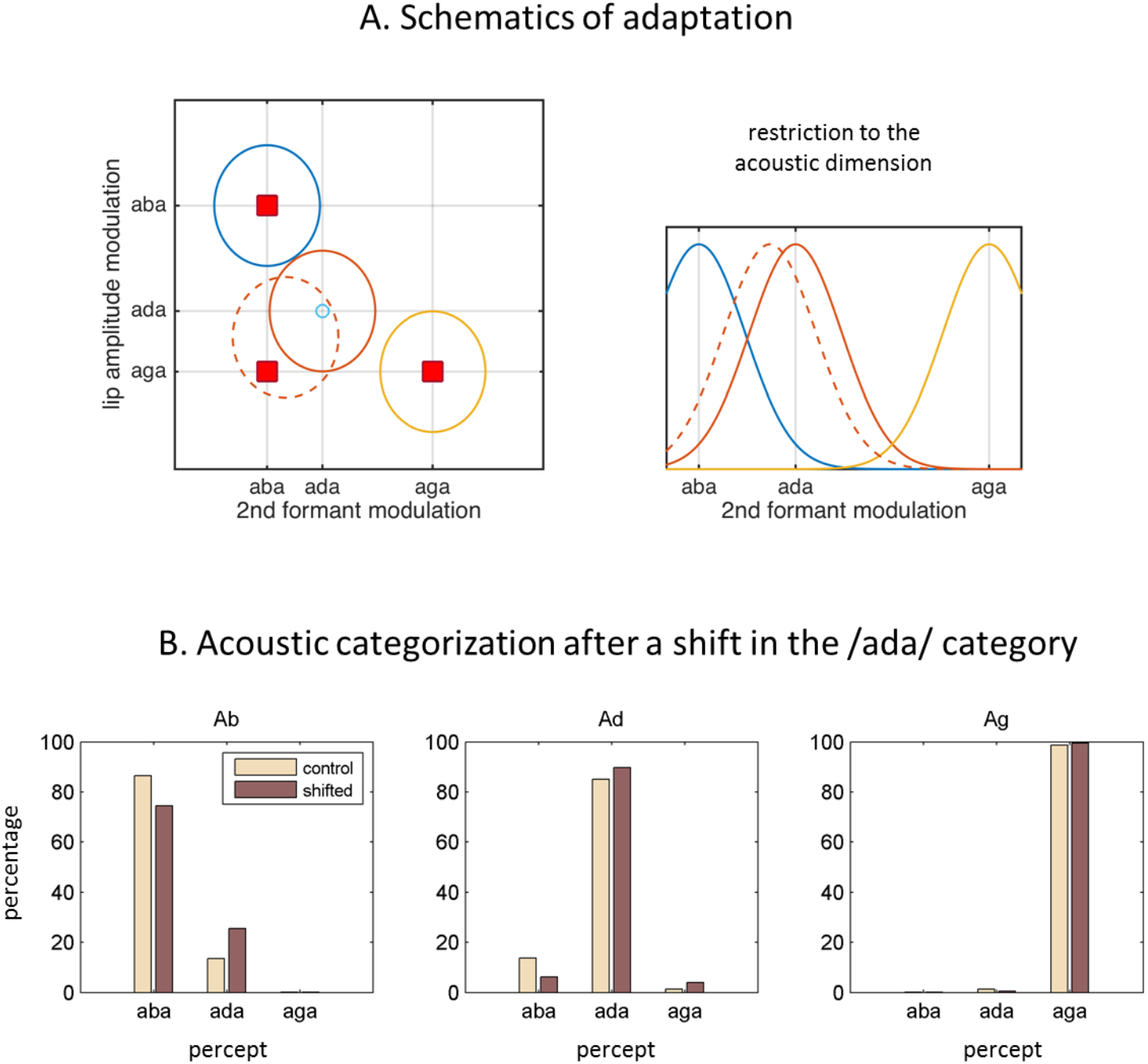
Internal model adaptation. A: Speech tokens are represented in a multimodal feature space here represented by two dimensions. Each ellipse stands for the internal representation of each congruent category (‘aba’ in blue, ‘ada’ in red, ‘aga’ in yellow). The red squares show the location of the audiovisual stimuli in the 2D feature space. They represent congruent /aba/ (top left), congruent /aga/ (bottom right), and McGurk stimuli (bottom left). When McGurk stimuli are repeatedly perceived as /ada/, the /ada/ representation (in solid red) is modified in such a way that it “moves” (dashed red) towards the presented McGurk stimulus (visual /aga/ with acoustic /aba/) and therefore should affect the processing of subsequent sensory input. The right panel shows the restriction to the acoustic dimension, illustrating how the acoustic representation for /ada/ has shifted towards that of /aba/. B: The effects of the shift in the internal representation on the categorization of the purely acoustic /aba/ (Ab), /ada/ (Ad) and /aga/ (Ag) sounds. Each panel shows the percentage of /aba/, /ada/ and /aga/ percepts for the “control” representations (solid lines) and the representations with the recalibrated /ada/ (dashed line). As in Lüttke et al. (2016), the biggest effect is observed in the categorization of the /aba/ sounds.

In the model, the residual prediction error occurs in the transformation from token identity to expected modulation of the acoustic feature (C_A_). Sensory evidence drives estimated C_A_ towards the experimentally presented value: /aba/ for McGurk stimuli. Thus, when the percept is /ada/, there is a mismatch between the top-down prediction as determined by the top-down component p(C|k) that drives C_A_ towards θ_/ada/,A_, and the actual value determined by the bottom-up component. This is evident in the following expression

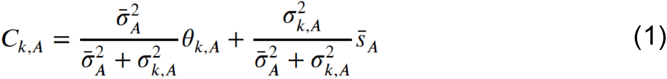

with the first term reflecting the prior expectation for category ‘k’ and the second reflecting the sensory evidence accumulated during the trial (see Eq 8 in the Materials and Methods section).

The expression can be rewritten to make the prediction error explicit.

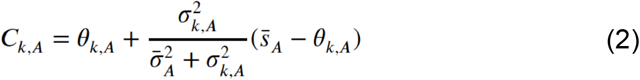

This expression highlights the prediction from the category (first term) and the residual prediction error in the second term.

To reduce the residual prediction error we consider that the participant recalibrates its generative model, that is, that it will change θ_k,v_ and θ_k,A_ driving them towards s_v_ and s_A_ respectively. If the stimuli are chosen with parameters “adapted” to the listener, as we do, s_V_ ~ θ_stim_,_V_ and s_A_ ~ θ_stim,A_.

We now present the results from running the experiment on 100 simulated listeners. A simulated listener is shown in Fig 2B. Each panel gives the histograms of responses (/aba/, /ada/ or /aga/) for each of the six types of stimulus. As in Lüttke et al. (Lüttke et al. 2016), accuracy for both acoustic and congruent audio-visual stimuli is high. In addition, there is a high frequency of fused percepts. We report the results obtained with different update rules; one derived from a Bayesian model as in (Kleinschmidt and Jaeger 2015), which assumes a stable environment, and two empirically motivated rules.

As a control we also simulated the experiment with no parameter updates (Fig 4A). The percentage of acoustic /aba/ categorized as /ada/ was 10% and the 95% CI for the difference between trials preceded or not by a fused McGurk stimulus was [-2.86 2.23]%.

**Fig 4.**
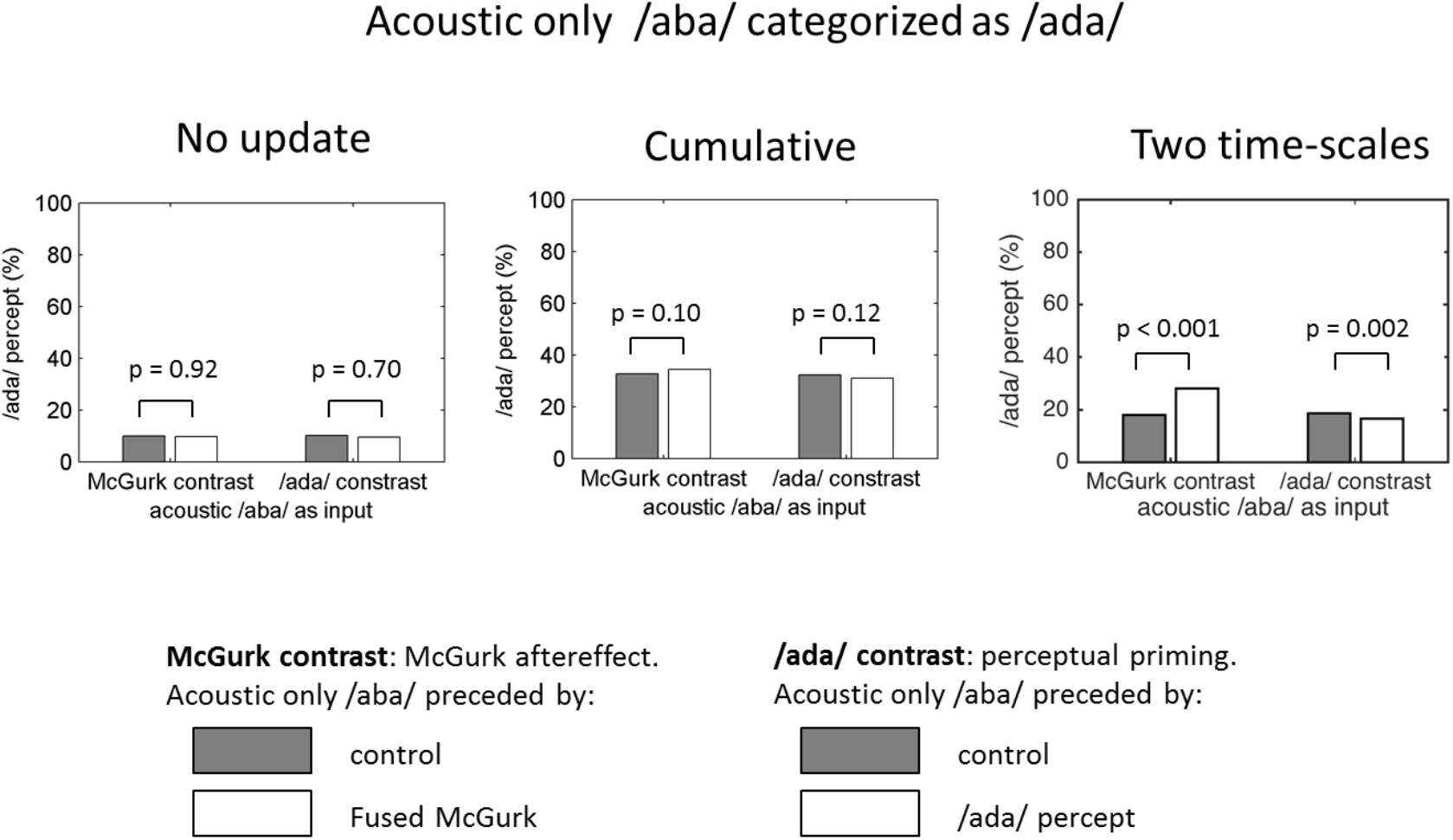
Cumulative and transient update rules. /ada/ percepts in response to acoustic /aba/ stimulation. We show two contrasts. The McGurk contrast compares the percentage of /ada/ responses when acoustic /aba/ is preceded by control stimuli (acoustic /aba/ and /aga/, congruent /aba/ and /aga/) versus fused McGurk trials. The /ada/ contrast refers to acoustic /aba/ preceded by control stimuli (acoustic /aba/ and /aga/) versus acoustic trials correctly perceived as /ada/. The three panels show the result of running the same perceptual model supplemented by three different update rules. Left, control, no update. Middle: constant delta rule. Right: update rule assuming that modification occurs at two time scales. The constant delta rule (middle) leads to changes in internal representations that are reflected in the overall increase of /ada/ percepts (with respect to the control, no update model on the left) however, it does not translate into effects on the previous trial. The model assuming two time scales does reproduce the effect of the fused McGurk on the next trial (McGurk contrast).

### Bayesian updating

We first considered the same kind of update rule used by Kleinschmidt and Jaeger (2015) to model changes in speech sound categorization after exposure to adapting stimuli. After each trial the generative model can update the parameters by considering the probability of the parameters given the sensory input and the categorization p(θ|k s_V_ s_A_), leading to sequential Bayesian updating (Eq 9 in the methods section; we run the simulations with κ_k,f,0_ = 3 and ν_k,f,0_ = 10). The percentage of /ada/ responses to acoustic only /aba/ stimuli was 17% both after fused McGurk and after all other control conditions. The 95% confidence interval for the difference in medians between the two conditions was [-2.35, 3.08]%, thus failing to reproduce the effect of interest. This might not be surprising; the Bayesian update rules have the form of a delta rule with a decreasing learning rate. As a consequence, the size of the changes in the categories diminishes as the experiment progresses and all stimuli end up experiencing the same internal model. The updates did lead to observable effects; there was an overall increase of /ada/ responses to acoustic /aba/ (17% vs. the 10% of the control experiment without parameter updates). The resulting changes in model parameters should lead to an after-effect, that is, the point of subjective equivalence in an /aba/-/ada/ acoustic continuum would be shifted in the direction of /aba/.

### Constant delta rule

The Bayesian update rule used above assumes that the parameters are constant in time and that therefore all samples have equal value, whether they are old or recent. This is equivalent to a delta rule with a learning rate tending to zero. We therefore considered a rule with a constant learning rate, which allows for updates of similar magnitude over the whole experiment. The model’s expected modulation for the perceived category was recalibrated according to:

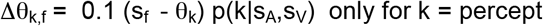

where *f* indexes the feature (*f*=*A*, acoustic feature; *f*=*V*, visual feature). As Fig 4B shows, the percentage of acoustic /aba/ categorized as /ada/ was not significantly higher when the preceding trial was a fused McGurk trial compared with any other stimulus (34% vs. 33%, median difference at 95% CI [-2.7 7.1]%). Note that in this case too, the proportion of /ada/ responses for acoustic /aba/ inputs is increased compared with simulations run without adaptation (Fig 4A).

Although the learning rate is constant, which means that recalibration magnitude does not necessarily decrease during the experiment, recalibration does not decay across trials. As a result, for trials between consecutive /ada/ percepts, stimuli experience the same /ada/ category and the simulations do not lead to a difference in how acoustic /aba/ is classified conditioned on whether acoustic /aba/ was preceded or not by a fused McGurk trial.

### Hierarchical Updates with intrinsic decay

We tested an alternative update rule based on the idea that it should reflect how changes occur in the environment. In this respect, we propose that one can characterize changes as occurring hierarchically, with just two levels in a first approximation. These would correspond to keeping “running averages” over different time scales, to be sensitive to fast changes without forgetting about longer-lasting trends.

We consider two sets of hierarchically related variables θ_k,f_ (fast) and μ_k,f_ (slow). The faster decaying one, θ_k,f_, is driven by both sensory prediction error and the more slowly changing variable, μ_k,f_ (more details can be found in Materials and Methods). This slowly changing and decaying variable, μ_k,f_, keeps a representation based on a longer term “average” over sensory evidence. In the limit, μ_k,f_ is constant; and this is what we consider here for illustrative purposes. Thus the update rules include a “jump” in the fast variable due to the sensory prediction error in the perceived category plus a decay term toward the slower variable for every category. The “jump” corresponds to the traditional update after an observation, the decay reflects the transient character of this update.

The results in Fig 4C were run with the following update rules:

Δθ_k,f_ = 0.1 (μ_k,f_ – θ_k,f_) p(k|s_A_,s_V_) for all k
Δθ_k,f_ = 0.3 (s_f_ – θ_k_) p(k|s_A_,s_V_) only for k = percept
Δμ_k,f_ = 0 for all k

where subscript f indexes the feature (f = A for acoustic, f = V for visual). All categories decay toward the corresponding long-term stable values (μ_k,A_, μ_k,V_) in the inter-trial interval.

The percentage of acoustic /aba/ categorized as /ada/ after control trials was 18% vs. 28% after fused McGurk, the median difference being 95% CI: [5.8 13.0]%. Therefore two effects can be observed; the overall increase in acoustic /aba/ categorized as /ada/ and the rapid recalibration effect reflected in the specific increase observed when acoustic /aba/ was preceded by a fused McGurk trial.

## Discussion

Speech sound categories are constantly revised as a function of the most recently presented stimuli. In this modelling work we provide a possible account for rapid and transient recalibration of speech sound categories in response to a changing sensory environment. In this account internal models of speech are recalibrated so that they better reflect the sensory environment. Our account suggests that recalibration occurs at least at two levels characterized by two different decaying timescales. We propose recalibration at a fast decaying/short-term level (on the order of seconds) that weighs recent versus past sensory evidence more strongly than a more slowly decaying/long-term level that keeps a longer trace of past sensory evidence. These elements are important because speech sounds have both variable and stable components: one needs to adapt to a new speaker without forgetting our past experience made from others.

Our model was motivated by the experimental results by Lüttke et al. (Lüttke et al. 2016) showing that the categorization of /aba/ sounds differs depending on whether the previous trial was a fused McGurk trial or any of the other five speech stimuli presented in the course of their experiment. Since the reported effect was distinct from other well-documented across-trial dependency effects, such as perceptual priming or selective adaptation, we sought to model it solely considering changes in perceived speech categories without external feedback. Like previous approaches, ours builds on the idea that the brain achieves perception by inverting a generative model (e.g., Rao and Ballard 1999; Knill and Pouget 2004; Karl J. Friston 2005) and by continuously monitoring the performance of this model to adapt it to the current stimulus landscape. One way the brain can change the model without external feedback is by using the perceptual outcome as a teaching signal. The latter informs the model so that the perceptual outcome better aligns with the sensory features leading to it (e.g. Kleinschmidt and Jaeger 2015). This agrees with results showing that humans seem to use their own categorical perceptual decisions as *ground truth* to inform a sensory identification task (Luu and Stocker 2018).

Recalibration of speech sound categories has been described within an “ideal adapter” framework (Kleinschmidt and Jaeger 2015). In this framework, speech sound categories are subject to trial-by-trial changes well described by a Bayesian approach that implicitly assumes that sound categories are stable within an experimental session, and hence uses update/learning rules that give the same weight to recent and past evidence. This makes sense in a stable environment where information remains equally relevant independent of its recency. While this adaptation rule successfully describes the dynamics of adaptation in sublexical speech categories in experiments with blocks of constant repeated stimuli (Bertelson, Vroomen, and De Gelder 2003; Vroomen et al. 2007), it failed here to account for the specific transient effect in Lüttke et al. (Lüttke et al. 2015).

Yet, we could successfully reproduce the results of Lüttke et al. (2016) by proposing empirically motivated update equations that include two hierarchically related variables with different learning and decaying timescales; a fast changing variable driven by current sensory prediction errors, and a more slowly changing one that acts as a longer-span buffer (Fig 5, equations (8) and (9)). As a result, the variable controlling the model expectations decays toward the longer-term variable after a transient change driven by the current sensory prediction error. Thus, after a fused McGurk trial, the acoustic representation of the /ada/ category was shifted towards that of acoustic /aba/ (Fig 3A), which increased the chances of an acoustic /aba/ to be categorized as /ada/ (Fig 3B). Since the shift decays over time, the increase in /ada/ percepts for acoustic /aba/ was most prominent when the acoustic /aba/ trial immediately followed the fused McGurk trial. On the other hand, since the decay goes towards a slower changing “version” of the /ada/ category, the model could also accommodate a more persistent accumulated adaptation as in Kleinschmidt and Jaeger (2015).

**Fig 5.**
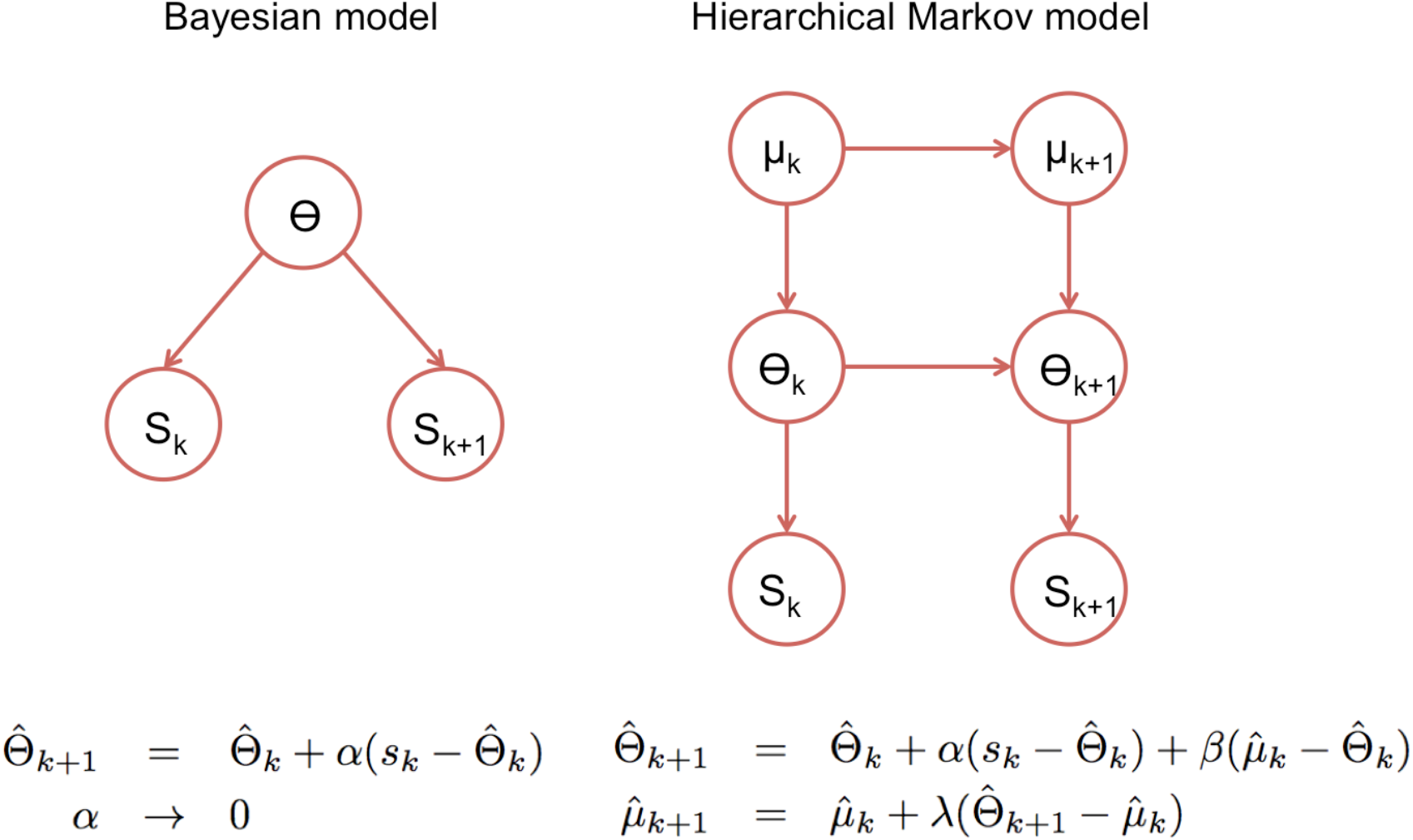
Statistical models underlying the two classes of update rules used in the paper. Here θ stands for the model parameters that determine the speech categories used by the perceptual model (See Fig 3) and ‘k’ for the trial index. On the standard Bayesian approach (left), model parameters are considered constant in time leading to update rules that give the same weight to all prediction errors, which in turn leads to a “learning rate” αk that becomes smaller with the number of trials. On the right we show a hierarchical Markov model implementation that would lead to the kind of update rules that we introduced empirically to accommodate the rapid recalibration effect. This alternative view implicitly assumes that model parameters can change in time and therefore lead to update rules with learning rates (α, β and λ) that under certain assumptions, settle to non-zero constant values.

In this work, we focused on the recalibration of the acoustic features associated with speech categories. However, given that lipreading can also be recalibrated by audiovisual input (Baart and Vroomen 2010), we would predict a corresponding effect on lipreading, namely that a purely visual /aga/ would be more readily categorized as /ada/ immediately after a fused McGurk trial. In summary, we propose that during audiovisual speech integration, and in McGurk stimuli in particular, the brain tries to find the most parsimonious explanation for the input even at the expense of residual prediction errors at the sensory periphery. Different participants may value differently this parsimony/accuracy trade-off; we hypothesize that those who consistently fuse the two streams change their templates for the /ada/ category, that is, for the perceived category, rather than for how the acoustic and visual features are processed in the periphery. The change in /ada/ effectively leads to category boundary shifts in acoustic continua, boundary shifts that are transient and will only add-up after consistent repetitions. Importantly, these effects are distinct from perceptual bias and specific adaptation, and from other documented short-term effects due to changes in the relative informational content of partially redundant acoustic features (Heald, Van Hedger, and Nusbaum 2017; Gabay and Holt 2018).

### Activity in auditory cortex

Lüttke et al. (2016) also noted that the behavioural effect was accompanied by distinct activity in auditory cortex. When an acoustic-only /aba/ was presented after a fused McGurk trial, fMRI activity in auditory cortex was more frequently classified as /ada/ by a learning algorithm trained to distinguish correctly identified acoustic /aba/, /ada/ and /aga/.

We have argued above that the intermediate level in our perceptual model could be interpreted as the representation of acoustic features in auditory cortex and visual features in extrastriate visual cortex. The acoustic feature in the model is the amplitude of the 2^nd^ formant transition: C_A_. As the perceptual process progresses within the trial, C_A_ is updated by moment-to-moment bottom-up prediction errors and by updated top-down predictions dominated by the /ada/ category in trials that will subsequently be categorized as /ada/ by the simulated participants. Since an /ada/ percept for acoustic /aba/ occurs more frequently after a fused McGurk trial, C_A_ will correspondingly be closer to C_A_(ada) than on trials preceded by the other types of stimuli, which lead to a lower proportion of /ada/ percepts. Therefore, our generative model, just as shown by the fMRI data, would predict activity in auditory cortex closer to /ada/ during the acoustic /aba/ after the fused McGurk trials, owing to the top-down input from the speech category level in the course of the within trial perceptual process.

### Revised “ideal adapter”

While the “ideal adapter” from Kleinschmidt and Jaeger (2015) focused on cumulative recalibration effects, our results suggest that shorter-lived effects are also behaviorally relevant. While the ideal adapter could be formalized as incremental optimal Bayesian inference in a non-volatile environment (Fig 5A), our empirically motivated update rule could possibly be cast in a normative framework by explicitly accounting for the volatility in the environment. True changes in the environment are sometimes referred to as ‘volatility’ to distinguish them from the trial-by-trial variability observable in a fixed environment. A hierarchical Markov model as the one shown in the right panel of Fig 5 could translate into hierarchical update equations of the form of the empirically proposed ones (Wilson, Nassar, and Gold 2013). This is because in a volatile environment current information is more relevant than past information, which leads to the notion of “optimal forgetting”. If the level of volatility is stable, one obtains a delta rule with a learning rate that converges onto a constant, with higher learning rates (faster “forgetting”) related to higher volatility levels. In Fig 5B it is assumed that the higher level (μ) has lower volatility than the intermediate level (θ). Therefore, our model combines volatility with hierarchy. This combination departs from models used to explain decision making in changing environments (Behrens et al. 2007; Summerfield, Behrens, and Koechlin 2011; C. D. Mathys et al. 2014), which are not hierarchical, and focus on the nontrivial task of inferring the volatility of the environment. The above studies provide evidence that human participants do adapt their learning rates when volatility is changed. Human performance in these tasks could be modelled without the need to keep representations over several time scales. It is important to note that in these tasks participants need to keep track of changes in arbitrary cue-reward associations or in arbitrarily defined sensory categories (Summerfield, Behrens, and Koechlin 2011). Our work refers to the adaptation of sublexical speech categories, that is, of already existing, learned and behaviourally relevant categories. The brain, therefore should not simply keep an estimate that changes flexibly to the local temporal environment, but probably benefits from also keeping a long-term estimate as an empirical “prior”.

Interestingly, hierarchically related variables with increasing time scales have been shown in a modelling study to increase the capacity of memory systems and improve the stability of synaptic modifications (Benna and Fusi 2016); and a model of synapses with a cascade of metastable states with increasing stability can flexibly learn under uncertainty (Iigaya 2016). The latter also modelled the intrinsic decay of synaptic modifications. The decay was faster for the more labile memory states, which are also those memory states that are more sensitive to new evidence. The deeper, more stable memory states showed slower decay. This maps nicely to our proposal.

Temporal hierarchies might also reflect the hierarchical nature of perception, in which higher level representations are increasingly abstract. If we see perception as a tower of abstractions, adaptation might work at every level of the hierarchy, with more abstract categories integrating update information within increasingly longer time windows, and therefore being increasingly stable.

Our modelling data are relevant in the domain of continuous speech processing, in particular to account for auditory processing anomalies in dyslexia. Evidence from a two-tone frequency discrimination task suggests that participants’ choices are driven not only by the tones presented at a given trial, but also by the recent history of tone frequencies in the experiment, with recent tones having more weight than earlier ones (Jaffe-Dax, Frenkel, and Ahissar 2017). It turns out that, when compared with controls, dyslexics show a decreased reliance on temporally distant tones, suggesting a shorter time constant (Jaffe-Dax et al., 2017). Translating this result to the current model, we could hypothesize that in dyslexia the long-term component (μ in Fig 5B) has either a shorter time span, or is coupled to the lower representation with a lower weight. In both cases, we would expect a deficit in building long-term stable speech category representations since they would be overly driven by the current context. ASD individuals on the other hand, are optimally biased by long-term tones, but do not show the bias by short-term tones of neurotypical participants (Lieder et al. 2019), which suggests a faster decay or an absence of the short-term component in Figure 5B. This would predict a failure of ASD individuals to show the specific effect after McGurk trials in the experiment simulated here.

### Conclusion

We have presented a revised “ideal adapter” model for speech sound recalibration that has both transient and cumulative components organized hierarchically. This new model allows us to interpret Lüttke et al.’s finding (2016) as evidence for a hierarchy of processes in the recalibration of speech categories. Our model further highlights that after experiencing the McGurk effect, it is not the acoustic features related to the sensory input (/aba/) that are modified, but the higher-level syllabic representation related to the percept (/ada/). This suggests that part of the sensory cortices activity changes that encode specific acoustic cues are not locally generated but reflect the interaction of bottom-up peripheral sensory inputs and top-down expectations from regions where categorical perception takes place. As far as speech perception is concerned, our results show that it could be fruitful to consider natural speech processing (including new speakers, languages, presence of noise etc…) as the inversion of a continuously monitored internal generative model adapted at different temporal scales closely reflecting its hierarchical structure.

## Materials and Methods

The goal of inference is to establish which is the speech token that gave rise to the incoming sensory input. For our present purposes we restrict ourselves to three tokens /aba/, /ada/ and /aga/ (as in (Lüttke et al. 2016)). Although several acoustic and visual features can distinguish between them, we choose to model the 2^nd^ formant transition, which is minimal for /aba/ but increases for /ada/ and /aga/, and the degree of lip closure, which is maximal for /aba/ and less prominent for /ada/ and /aga/ (lip closure /aba/>/ada/>/aga/). This choice is based on the fact that what distinguishes between the three speech sounds is the place of articulation. Acoustically the 2^nd^ formant transition is an important cue for place of articulation, particularly within the ‘a’ vowel context (Liberman et al. 1957); visually it is the degree of lip aperture at the time of the vocal cavity occlusion depending on its location (complete lip closure for the bilabial (/aba/), and decreasing lip closure for the alveolar (/ada/) and velar /aga/) (Campbell 2008; Varnet et al. 2013).

The generative model has three levels; the higher level encodes the speech token, the speech token in turn determines the expected values for the audiovisual cues, as represented in Fig 1A. The model includes the three possible tokens, each determining the expected distribution of its associated audiovisual features. We also introduce uncorrelated noise to account for sensory variability. The parameters, location and spread of features associated with each token, as well as the level of sensory noise, define an individual listener’s internal model.

Although not strictly necessary for our present purpose, the model includes the temporal profiles of the two sensory features (Fig 1B). The latter are fixed and only their amplitude is modelled as a random variable that changes only across trials. The inclusion of the temporal dimension allows us to gain intuition into the separation of the perceptual and recalibration processes.

We use ‘k’ as the speech token index k = {/aba/, /ada/, /aga/}; ‘f as feature index f = {V,A}, where ‘V’ stands for the visual feature (lip aperture) and ‘A’ for the acoustic feature (2^nd^ formant transition). The idea is that the lip aperture and 2^nd^ formant have stereotypical temporal modulation profiles that vary in amplitude according to the speech token.

We use ‘M_f,n_’ to denote the temporal profile of the modulation for feature ‘f at time point ‘n’ (M_V,1_ … M_V,N_) stands for the temporal profile of lip modulation and (M_A,1_ … M_A,N_) for the 2^nd^ formant modulation profile. The time dependent modulation for each feature, lip ‘y_V,n_ :(y_V,1_ … y_V,N_)’ and 2^nd^ formant ‘y_A,n_ :(y_A,1_ … y_A,N_)’ is thus modelled as the product of a constant modulation amplitude ‘C_V_’ or ‘C_A_’ and a standard temporal profile ((M_V,1_ … M_V,N_) or (M_A,1_ … M_A,N_)). In this way, we are explicitly separating ‘what’ and ‘when’. Thus ‘y_f,n_’ represents feature ‘f at time point ‘n’. The full generative model linking a token ‘k’ to audiovisual sensory features is described by the equations below.

There are two sources of variability, moment-to-moment uncorrelated sensory noise (σ_V_, σ_A_) and the variability of modulation amplitudes associated with the same token in different realisations (σk,V, σk,A) k = {/aba/, /ada/, /aga/}.

The hierarchical generative model is defined by the following relations:

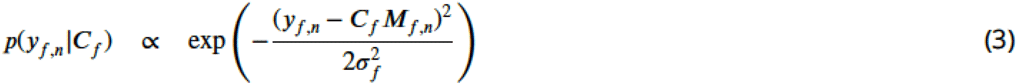

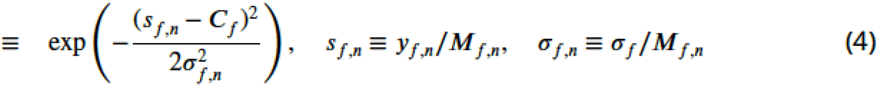

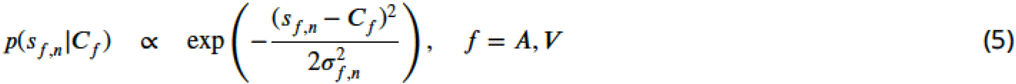

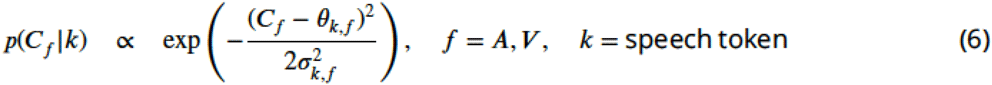

In the following we will work with ‘s_f,n_’ instead of ‘y_f,n_’, that is, with the information about the – constant-amplitude instead of the time varying feature profile. This makes the associated variance, (σ_f,n_)^2^, dependent on the standard temporal profile of the relevant feature.

While the above defines the generative (top-down model) p(s_f,n_|C_f,k_), our interest lies in its inversion p(k|s_f,1_, …, s_f,N_), where s_f,1_, …, s_f,N_ represents the sensory input in a single trial.

From the above equations one obtains:

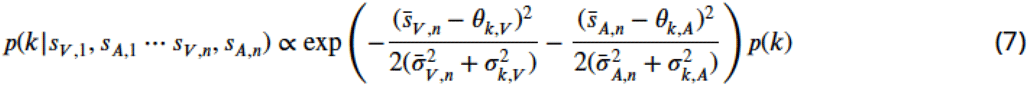

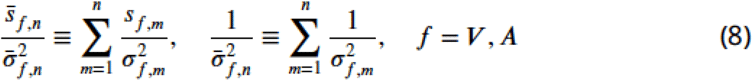

which results from marginalizing over ‘C_V_ and ‘C_A_’-the intermediate stages that encode the visual and acoustic features, explicitly:

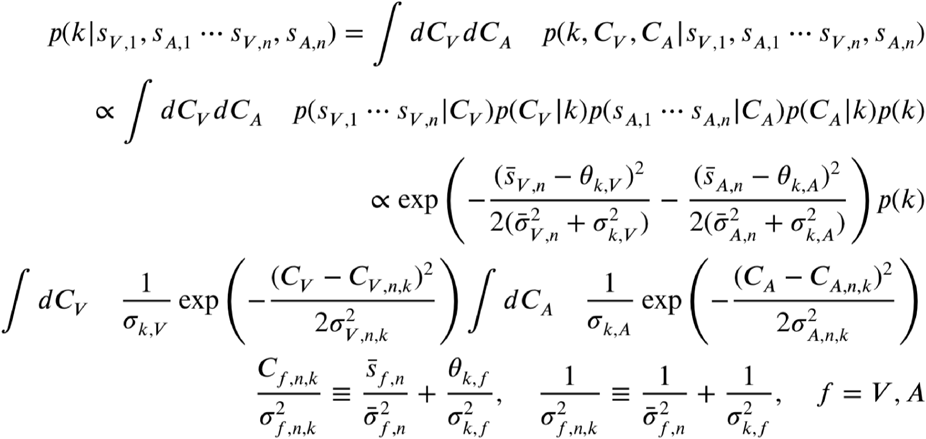

We assume that initial variance and prior probabilities are equal across categories (p(k)=1/3).

Alternatively, marginalization over ‘k’ gives the probabilities over the hidden variables ‘C_V_’ and ‘C_A_, which we associate with encoding of stimulus features (lip aperture, 2^nd^ formant transition) in visual and auditory cortex respectively.

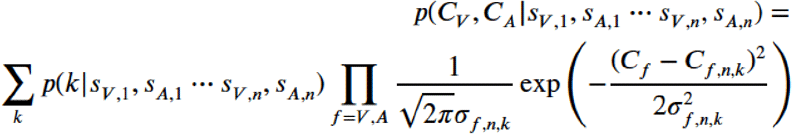

This shows explicitly how internal estimates of sensory features (lip aperture ‘C_V_’ and 2^nd^ formant ‘C_A_) are driven by bottom up sensory evidence (s_V_, s_A_) and top-down empirical priors, with each category ‘k’ contributing according to its internal expectations (θ_k,V_ θ_k,A_). When there is strong evidence for a given category ‘k’, the sum can be approximated by a single Gaussian centred at a compromise between θ_k,V_ and s_V_ for the visual feature and between θ_k,A_ and s_A_ for the acoustic feature.

The centre and variance of each Gaussian can also be written in terms of moment-to-moment incremental updates

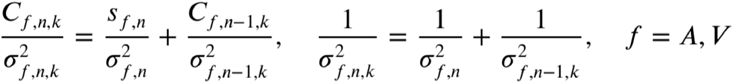

where *n* indexes the time point within the trial and *k* denotes the speech token. Or equivalently for the *C_fn,k_*:

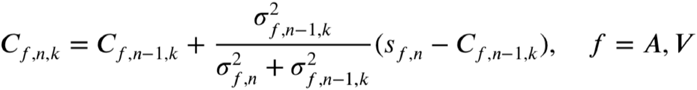

which explicitly shows how the internal representation of the modulation magnitudes is updated by moment-to-moment prediction errors.

In principle the model could also be made to perform causal inference (Magnotti and Beauchamp 2017), that is, decide whether the two sensory streams belong to the same source and therefore should be integrated, or whether the two sensory streams do not belong to the same source, in which case the participant should ignore the visual stream. Since Lüttke et al. explicitly selected the participants that consistently reported /ada/ for the McGurk stimulus, these subjects were fusing the two streams. We hence assume that integration is happening at every audio-visual trial.

The model’s percept corresponds to the category that maximizes the posterior distribution p(k|s_f,1_, …, s_f,N_).

### Recalibration model

The previous section presented how the model does inference in a single trial. We now turn to how the model updates the parameters that encode the internal representation of the three speech categories. This happens after every trial, thus simulating an internal model that continuously tries to minimize the difference between its predictions and the actual observations; we assume that in this process, in which the model tries to make itself more consistent with the input just received, it will only update the category corresponding to its choice. We will present three updating rules. The normative incremental Bayesian update model used by (Kleinschmidt and Jaeger 2015), and the empirically motivated constant delta rule and hierarchical delta rule with intrinsic decay.

### Bayesian updating

The internal representation of the speech categories is determined by six location parameters (θ_k,V_ θ_k,A_) and six width parameters (σ_k,V_, σ_k,A_). We follow Gelman et al. (Gelman et al. 2003) and define the following prior distributions for the internal model parameters (θ_k,V_, σ_k,V_; θ_k,A_, σ_k,A_). For each of the six (θ, σ) pairs (2 sensory features × 3 categories) the prior is written as:

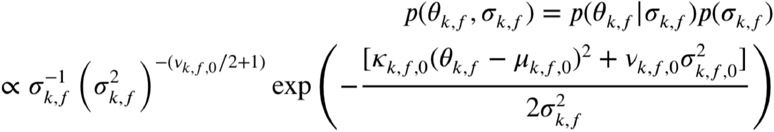

As above, f refers to the sensory feature, either V or A, and k to the speech category, either /aba/, /ada/ or /aga/.

After a new trial with sensory input (s_V_, s_A_) the updated prior has the following parameters:

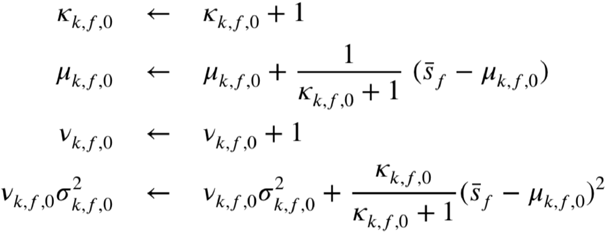

After each trial the inference process described in the previous section determines the percept from the posterior probability p(k|s_V_ s_A_). Only the feature parameters of the representation corresponding to the percept are subsequently updated.

We use the values that maximize the posterior over the parameters given the input and the current estimated category ‘k’ to determine the point estimates that will define the updated model parameters for the next trial (Eq 6). The updates for the location and spread parameters then take the form:

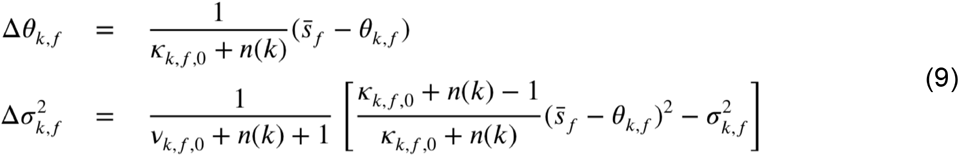

Where ‘k’ is the perceived category, n(k) the number of times the category has been perceived, f =V,A designates the sensory feature and ν_k,f,0_ and κ_k,f,0_ are parameters from the prior distribution. The larger ν_k,f,0_ and κ_k,f,0_ are, the more k “perceptions” it takes for the parameter values of category ‘k to plateau but also the smaller the updates after each trial.

### Constant delta rule

The above update equations implicitly assume that the environment is stable and therefore updated parameters keep information from all previous trials. This is the result of the generative model, which did not include a model for environmental parameter changes. Introducing expectations about environmental changes led us to consider rules with constant learning rates. We restrict ourselves here to updates for the six location parameters (θ_k,V_ θ_k,A_) of Eq 6.

We first considered a constant delta rule scaled by the evidence in favour of the selected category p(k|s_V_, s_A_),

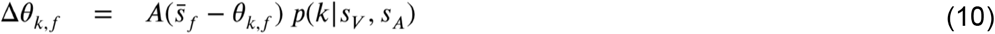

As in the Bayesian case, updates accumulate without decay between trials. The main difference is that θ_k,f_ is driven more strongly by recent evidence than by past evidence, implicitly acknowledging the presence of volatility.

### Hierarchical delta rule with decay

Finally we consider updates that decay with time. We reasoned that the decay should be towards parameter estimates that are more stable, which we denote by μ_k,f_. We propose a hierarchical relation, with updates in μ_k,f_ being driven by θ_k,f_, while updates in θ_k,f_ are driven by sensory evidence. At each trial all categories (k’) decay toward their long-term estimates and only the perceived category (k) updates both μ_k,f_ and θ_k,f_ :

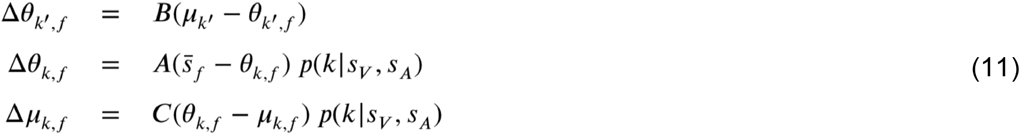

The first equation reflects the decay while the last two equations apply to the perceived category ‘k’. ‘A’, ‘B’ and ‘C’ are constant parameters.

### Model simulations

We simulate the experimental paradigm in Lüttke et al. (Lüttke et al. 2016), in which human participants were asked about what they heard when presented with auditory syllables or auditory syllables accompanied with a video of the corresponding speaker’s lip movements. There were 6 stimulus types: three acoustic only stimuli: /aba/, /ada/ and /aga/ and three audiovisual stimuli, congruent /aba/, congruent /ada/ and McGurk stimuli, that is, acoustic /aba/ accompanied by the video of a speaker articulating /aga/. In the original experiment three different realizations of each of the 6 types were used. In our simulations we use a single realization per stimulus.

Our model simulates a single participant. As described above it proposes that syllables are encoded in terms of the expected amplitudes and variances of audiovisual features. The expected amplitudes were taken from the mean values across 10 productions from a single male speaker (Olasagasti, Bouton, and Giraud 2015), the amplitudes were then normalized by dividing by the highest value for each feature. The remaining parameters in the model, variances and sensory noise levels, were chosen so that the overall categorization results, percentage of /aba/, /ada/ and /aga/ responses to the 6 types of stimuli were qualitatively similar to those reported by Lüttke and colleagues. There was no explicit effort to fit the model, since our goal is to present a plausibility argument. What we asked was that acoustic and congruent audiovisual items were mostly correctly categorized and that McGurk stimuli were mostly categorized as /ada/, thus fulfilling the conditions of the participants chosen for the analysis in (Lüttke et al. 2016).

The model parameter values are given in Table 1.

**Table 1.** Model parameters.

|  |  |  |  |  |
| --- | --- | --- | --- | --- |
| $\theta_{/aba/,A} = 0.1$ | $\theta_{/ada/,A} = 0.4$ | $\theta_{/aga/,A} = 1$ | $\sigma_{k,C} = 0.1$ | $\sigma_A = 0.12$ |
| $\theta_{/aba/,V} = 1$ | $\theta_{/ada/,V} = 0.6$ | $\theta_{/aga/,V} = 0.37$ | $\sigma_{k,C} = 0.1$ | $\sigma_V = 0.06$ |

The first three columns correspond to the initial location parameters of the three categories along the acoustic and visual feature dimensions. The 4<sup>th</sup> column gives the initial standard deviations of the three categories. The last column gives the values of sensory noise in the two modalities.

The stimuli presented to the modelled participant corresponded to the expected acoustic and visual features of their internal model. Thus if the internal model for /aba/ is centered at θ_/aba/A_ for the acoustic feature and at θ_/aba/V_ for the visual feature, those are the amplitudes chosen for the input stimuli. In other words, stimuli were tailored to the modelled participant. The six stimuli were defined by:

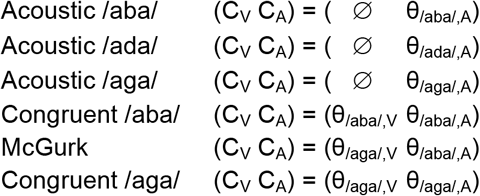

Even if the underlying parameters for a given stimulus type were the same for every trial, sensory noise created variability. The input to the model was the pair of temporal sequences y_V_(t) and y_A_(t) defined by:

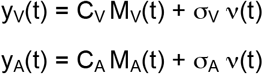

The lip and 2^nd^ formant temporal modulation profiles (M_V_(t) and M_A_(t), Fig 1B) were defined as in (Olasagasti, Bouton, and Giraud 2015). Each profile, representing the intervocalic transition between the two vowels in the “a” vocalic context, was modelled with 289 time points. The profiles are normalized so that the sum of M^2^_A,n_ over ‘n’ (number of time points) is one. The last term represents independent Gaussian noise different for the two modalities.

The six stimulus types were presented to the model in random order. At the end of the presentation, the model chose a percept based on the posterior distribution over syllable identity ‘k’ given the stimulus. The perceived syllable was then recalibrated by updating its defining parameters (either both mean and variance or mean alone).

We looked at acoustic only /aba/ trials and compared the proportion of /ada/ responses in two “contrasts” as in Lüttke et al. (Lüttke et al. 2016). The McGurk contrast compares the proportion of /ada/ responses when the preceding trial was a fused McGurk trial versus other stimuli (acoustic /aba/ and /aga/ and congruent /aba/ and /aga/). The “ada” contrast looks at acoustic /aba/ trials preceded by correctly categorized acoustic /ada/ versus other stimuli (acoustic /aba/ and /ada/).

All simulations were performed using custom scripts written in MATLAB (Release R2014b, The MathWorks, Inc., Natick, Massachusetts, United States). Pair-wise comparisons were performed using MATLAB’s implementation of the Wilcoxon signed rank test.

While the original experiment had 27 participants and each stimulus type was presented 69 times, we run the experiment 100 times with 200 repetitions per stimulus type. The larger number of repetitions per experiment allowed a reduction in the sampling variability in the percentages of acoustic /aba/ categorized as /ada/. 95% confidence intervals were estimated by randomly sampling 27 simulations from the 100 available. For each simulation, the median difference between the proportion of acoustic /aba/ categorized as /ada/ after fused McGurk or other trials was calculated. The sampling described above was used to estimate the 95% confidence interval of the increase in proportion of /ada/ percepts after a fused McGurk trial.

